# Introduction of Barnase/Barstar in soybean produces a rescuable male sterility system for hybrid breeding

**DOI:** 10.1101/2023.04.24.538080

**Authors:** Nicole Szeluga, Patricia Baldrich, Ryan DelPercio, Blake C. Meyers, Margaret H. Frank

**Affiliations:** Plant Biology Section, School of Integrative Plant Science, Cornell University, Ithaca, NY, USA; Donald Danforth Plant Science Center, St. Louis. MO, USA

**Keywords:** Barnase, Barstar, male sterility, hybrid breeding, soybean, *Glycine Max*, tapetum

## Abstract

Hybrid breeding for increased vigor has been used for over a century to boost agricultural outputs without requiring higher inputs. While this approach has led to some of the most substantial gains in crop productivity, breeding barriers have fundamentally limited soybean (*Glycine max*) from reaping the benefits of hybrid vigor. Soybean makes inconspicuous flowers that self-fertilize before opening, and thus are not readily amenable to outcrossing. In this study, we demonstrate that the Barnase/Barstar male sterility/male rescue system can be used in soybean to produce hybrid seeds. By expressing the cytotoxic ribonuclease, Barnase, under a tapetum-specific promoter in soybean anthers, we are able to completely block pollen maturation, creating male-sterile plants. We show that fertility can be rescued in the F1 generation of these Barnase-expressing lines when they are crossed with pollen from plants that express the Barnase inhibitor, Barstar. Importantly, we found that the successful rescue of male fertility is dependent on the relative dosage of Barnase and Barstar. When Barnase and Barstar were expressed under the same tapetum-specific promoter, the F1 offspring remained male-sterile. When we expressed Barstar under a relatively stronger promoter than Barnase, we were able to achieve a successful rescue of male fertility in the F1 generation. This work demonstrates the successful implementation of a biotechnology approach to produce fertile hybrid offspring in soybean.

## Introduction

Soybean *(Glycine max)* is one of the most economically and societally impactful crops in the world, providing a significant percentage of all protein for animal consumption on a global scale, and playing key roles in oil production, manufacturing, and biofuel applications (American Soybean Association, n.d.). In 2022, 86.3 million acres of soybean were planted in the U.S., making soybean the second most planted crop after corn, which covered 93 million acres (Ates and Bukowski 2023a; Dohlman and Hansen 2022; American Soybean Association, n.d.). To keep up with the growing demand for soy-based animal feed, the USDA projects soybean acreage will increase by 19.6% by 2032 (Dohlman and Hansen 2022). Given the importance of soybean to global agriculture, advances in soybean productivity could have a transformative impact, and promote sustainable agriculture by enabling farmers to produce higher yields on existing acreage.

Hybrid vigor, also known as heterosis, is the phenomenon in which offspring outperform both of their parents and has been used for over a century to increase crop yields, improve abiotic and biotic stress tolerance, and enhance the nutritional quality of seeds (Fu et al. 2014; Lippman and Zamir 2007). Unlike crops that have benefited from hybrid vigor, such as maize, soybean has yet to reap the benefits of a large hybrid breeding program. Based on previous small-scale trials, soybean hybrids made through laborious hand-crossing techniques can produce a 10-20% increase in yield relative to inbred parents (Perez, Cianzio, and Palmer 2009; Burton and Brownie 2006; Palmer et al. 2001). Due to the challenging nature of producing soybean hybrids, these trials are limited to a small number of genotype combinations, indicating that the potential for yield improvement through heterosis is far from exhausted. To capture the benefits of hybrid breeding, the production of high-yielding and commercially available hybrid soybean seeds must be efficient, affordable, and compatible with existing farming practices.

One approach to increase outcrossing without emasculating flowers is to block self-fertilization using male sterility (MS). The combination of nuclear male sterility (NMS), cytoplasmic male sterility (CMS), and photoperiod/temperature-sensitive genic male sterility systems with fertility restorer genes have produced promising results in small trials of soybean hybrids (Nadeem et al. 2021; Ramlal et al. 2022; Palmer 2000; Thu et al. 2019; Palmer et al. 2001). However, multiple drawbacks exist in these systems, making them near impractical for commercial hybrid soybean breeding applications. For example, the three-line breeding system requires a maintainer line, results in low genetic diversity occurring from inbreeding CMS lines, and requires restrictive growth conditions in the case of photoperiod/temperature-sensitive male sterility systems (Nadeem et al. 2021; Li et al. 2019; Bai and Gai 2006). Most importantly, many of these male-sterile lines are not 100% effective at blocking self-fertilization and often fail to exhibit a full rescue of fertility in the F1 generation, making these existing solutions for hybrid breeding commercially inviable (Bai and Gai 2006).

An alternative to NMS and CMS is the combination of Barnase and Barstar; a two-component system in which obligate outcrossing plants are created through the expression of a tapetum-specific cytotoxic ribonuclease (Barnase) and its inhibitor (Barstar) (Hartley 1989, 1988, 2001). In wildtype post-meiotic anthers, each pollen sac is composed of a ring of endothecium cells, a middle layer, and a tapetal layer that surrounds developing pollen grains. The role of the tapetum is to provide nutrients and precursory material to build the outer pollen coat for developing grains. Targeted expression of Barnase in the tapetum destroys this cell layer, causing male pollen infertility (Mariani et al. 1990, 1992). Fertility is restored in the second generation by crossing Barnase-expressing plants with pollen carrying a transgene for the tapetum-specific expression of the ribonuclease inhibitor, Barstar (Mariani et al. 1992; Hartley 1989, 2001, 1988; Mariani et al. 1990). This approach has been adopted to create hybrid breeding systems for several crops, including a variety of vegetables and oil seed crops (Jagannath et al. 2002; Mariani et al. 1992; Colombo and Galmarini 2017).

In this study, we integrate Barnase/Barstar male sterility/male rescue technology into soybean to test the potential for this biotechnological approach to enable large-scale hybrid breeding in soybean. We generated Barnase expressing lines that consistently exhibited complete male sterility and two different rescue lines that expressed Barstar at equal or higher levels than Barnase. We show that a higher dose of Barstar relative to Barnase is essential for producing a successful rescue of male fertility in the hybrid generation of Barnase male sterility lines. This work opens up new resources for investigating heterosis and advancing the future of breeding in soybean.

## Results

### Designing tapetum-specific promoters to develop a rescuable male sterility system in soybean

To isolate the cytotoxic ribonuclease activity of Barnase and rescue activity of Barstar to the tapetal cell layer of soybean anthers, we identified two soybean paralogs for the previously characterized tapetum-specific Arabidopsis gene TA29 (Atg07230) (Mariani et al. 1990, 1992). TA29a (Glyma.09G144000) and TA29b (Glyma.16G197100) are specifically expressed in unopened soybean flowers with TA29a exhibiting higher expression than TA29b (Figure S1) (Sreedasyam et al. 2022; Valliyodan et al. 2019). To confirm that TA29a/TA29b are specifically expressed in tapetal cells, we generated a dual reporter line with TA29a::tdTomato and TA29b::ZsGreen (Figure 1d). In this reporter line, we observed tapetum-specific expression of both fluorophores in the early stages of microspore development (Figure 1d-o), indicating that these TA29 paralogs are appropriate for directing Barnase and Barstar expression in soybean.

**Figure 1:**
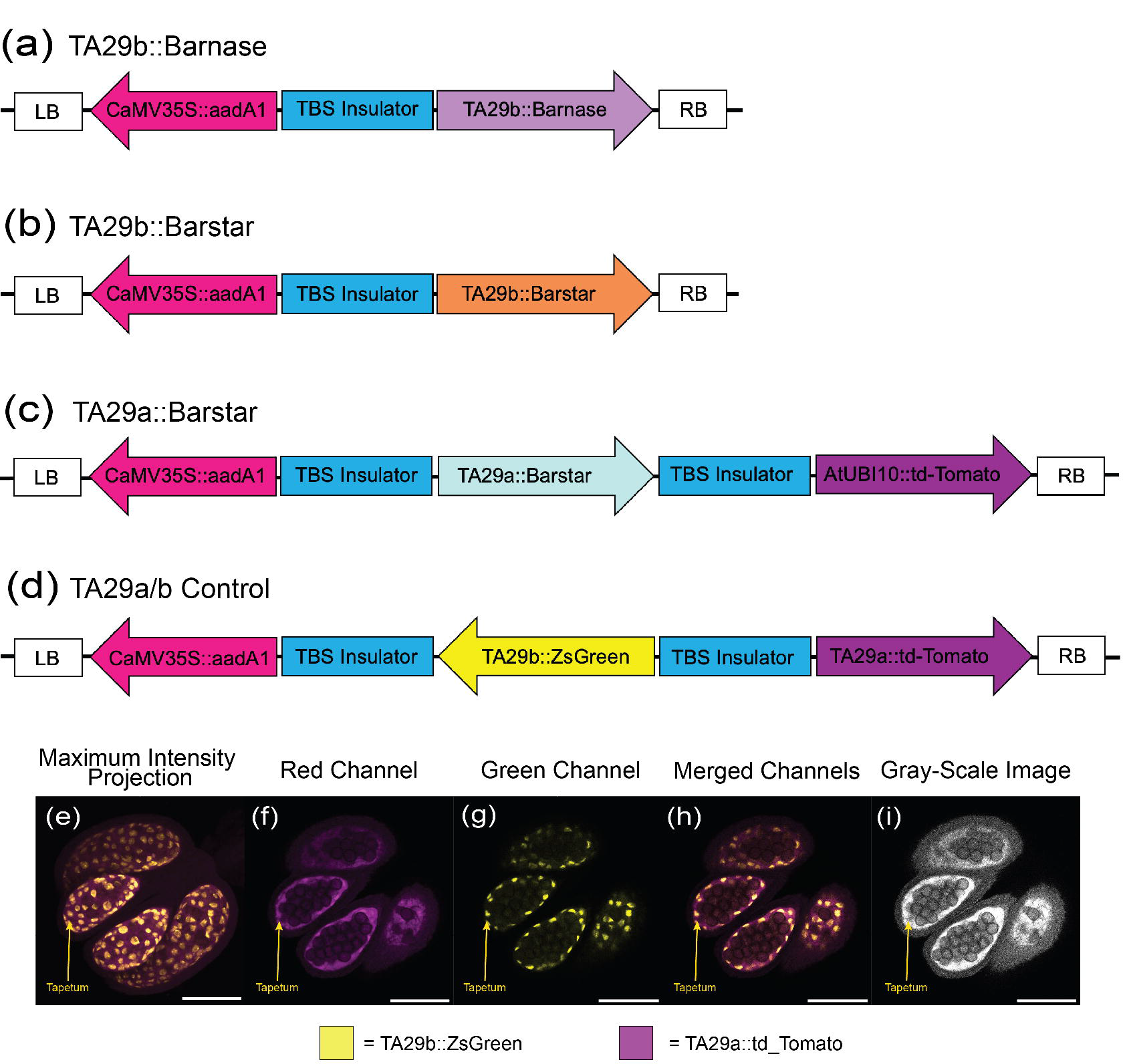
Construct designs for tapetum-specific expression of Barnase and Barstar generates stable and spatially accurate gene expression in soybean anthers. (a,b) Tapetum-specific expression of Barnase and Barstar driven by the weaker promoter, TA29b. aadA1 is a resistance gene for kanamycin selection. (c) Construct design for Barstar expression under stronger promoter, TA29a. This construct design also includes constitutive expression of the fluorophore, td_Tomato. (d) Construct design for TA29a/b control line. Anther images of the TA29a/b control line (e-i) were cleared and imaged with confocal microscopy. Red and Green Channels were altered in the Fiji software system to project magenta and yellow, respectively. The white scale bar is 100μm.

Since Barnase is a potent ribonuclease, we decided to take advantage of the differential expression of TA29a/TA29b to create a rescue system in which Barstar dosage is consistently higher than Barnase. We hypothesized that this differential dosage would limit the presence of uninhibited Barnase which could negatively impact male fertility in the F1 rescues. To test this hypothesis, we designed one TA29b::Barnase expressing construct, and two TA29a::Barstar and TA29b::Barstar rescue constructs. In addition, we tagged TA29a::Barstar with an AtUBI10::td-Tomato fluorophore to facilitate quick identification of successful crosses onto TA29b::Barnase pre-genotyping (Figure 1a-c).

### Tapetum-specific expression of Barnase produces complete male sterility

Of the 13 independent TA29b::Barnase lines that we generated, 12 exhibited consistent male sterility phenotypes (Figure S2). Wildtype soybean plants produce both cleistogamous (closed) and chasmogamous (open) flowers at maturity. Interestingly, while there were no other alterations to overall flower morphology and size, we noticed that all 12 of the male-sterile Barnase lines only produced cleistogamous flowers (Figure 2g). To evaluate the effect of Barnase expression on stamen and carpel formation, we dissected and imaged flowers at peak reproductive maturity (Figure 2g-k). All of the TA29b::Barnase male-sterile lines formed anthers that were misshapen, either opaque white/light yellow or semi-translucent in appearance, and failed to form pollen grains (Figure 2h-k, Figure S2b,e,h,k). In contrast, wildtype anthers were golden and shedding pollen at maturity (Figure 2b,c). We used propidium iodide (PI) staining and confocal imaging to further investigate the impact of Barnase on internal anther organization and identify the cellular basis for male sterility in these lines. In WT post-meiotic anthers, we observed pollen grain release from mature pollen sacs, whereas, in Barnase-expressing plants, the cavity of the anther lacked microspores and instead exhibited degenerating cellular debris that fluoresced brightly with PI staining (Figure 2c,i).

**Figure 2:**
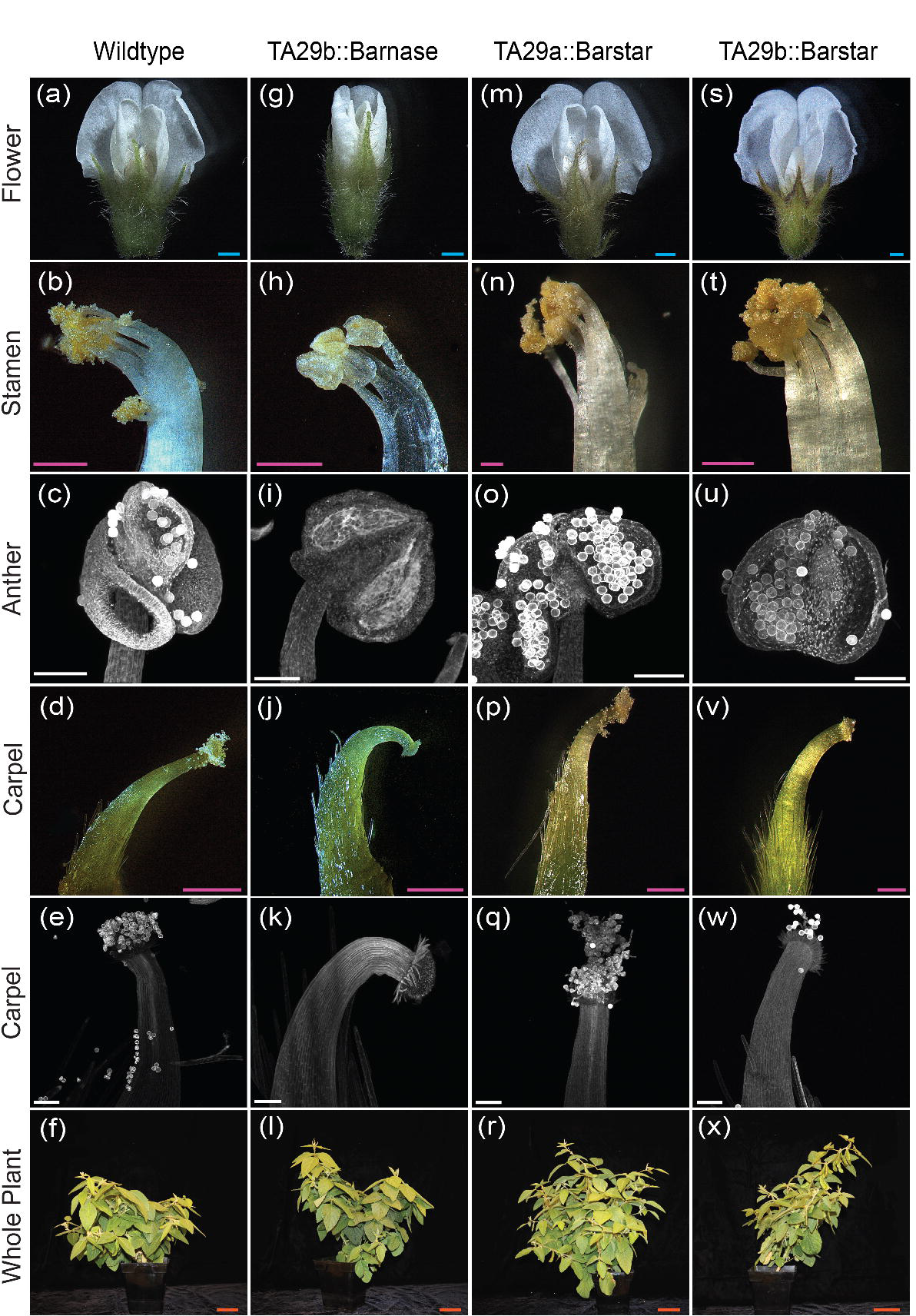
Flower dissection of TA29b::Barnase displays cleistogamous flowers, male-sterile anthers, and curved carpels compared to wildtype, TA29a::Barstar, and TA29b::Barstar. Flowers (a,g,m,s), stamen (b,h,n,t), and carpel images (d,j,p,v) were dissected and imaged under a dissecting microscope. Whole flower images were captured at peak maturity. Anther (c,i,o,p) and carpel images (e,k,q,w) were stained with PI and imaged using confocal microscopy. Soybean plants (f,l,r,x) were grown under growth chamber conditions and imaged 90 days post-germination. Scale bars for the figure are as follows: the blue bar is 1mm; the pink bar is 500μm; the white bar is 100μm; the orange bar is 100mm.

To investigate whether TA29b::Barnase impacted carpel formation, we imaged overall carpel shape and papillar cell morphology using a dissecting microscope and confocal imaging, respectively. When compared to WT, we noticed that mature carpels in Barnase expressing plants tended to curve inward, positioning the stigma towards the center of the flower where the free stamen is located (Figure 2d,e,j,k) (Talukdar and Shivakumar 2012; Singh, Chung, and Nelson 2007). We found this phenotype to be consistent across independent TA29b::Barnase transformants (Figure S2c,f,i,l). WT carpels also exhibited slight curvature, however, the stigma tended to face upward in these flowers. Congruent with our findings that the Barnase plants were completely male-sterile, we found zero fertilized pods on any of these lines (Figure S3). Interestingly, in lieu of fertilized pods, we did observe parthenocarpic pod formation on the TA29b::Barnase (Figure S3). These stubby pods were formed as a result of ovule development in the absence of fertilization, and could easily be distinguished from viable pods based on their length, presence of unfertilized ovules in place of seeds, and greatly reduced size (Figure S3).

Since Barstar is an inhibitor of Barnase, we did not expect TA29a::Barstar and TA29b::Barstar to differ in phenotype from WT. In line with this hypothesis, the petals, carpels, and stamens for these two transgenic lines were indistinguishable from those formed on WT flowers (Figure 2a-e, m-q, s-w). Furthermore, the two Barstar lines produced numerous fertile pods similar to the WT controls. To test whether Barnase and Barstar constructs impacted the overall phenotype of WT, we collected a spectrum of image-based measurements that relate to vegetative health and photosynthetic capacity using a CropReporter™ system (Netherlands Plant Ecophenotyping Center) and corresponding Phenovation Data Analysis software v548. We did not identify any noticeable change in the vegetative health of either the Barnase or Barstar transgenic lines relative to WT plants (Figure S4). This data demonstrates that the Barstar constructs have no measurable impact on plant phenotype, and the Barnase construct primarily impacts male reproductive development.

### Rescuing male fertility to generate F1 hybrids

To test whether Barnase-expressing carpels are receptive to pollen, we crossed WT and TA29a::Barstar pollen onto TA29b::Barnase stigmas and used aniline blue staining to track pollen tube elongation. Relative to the WT control, we observed unaltered pollen grain germination and growth on the TA29b::Barnase carpels (Figure 3a-c). Our confocal images show that pollen grains were able to adhere to the surface of the stigma, and regardless of the pollen and/or carpel genotypes, we observed comparable levels of pollen tube growth at 24 hours post-pollination (Figure 3a-c). This data demonstrates that TA29b::Barnase carpels are reproductively viable.

**Figure 3:**
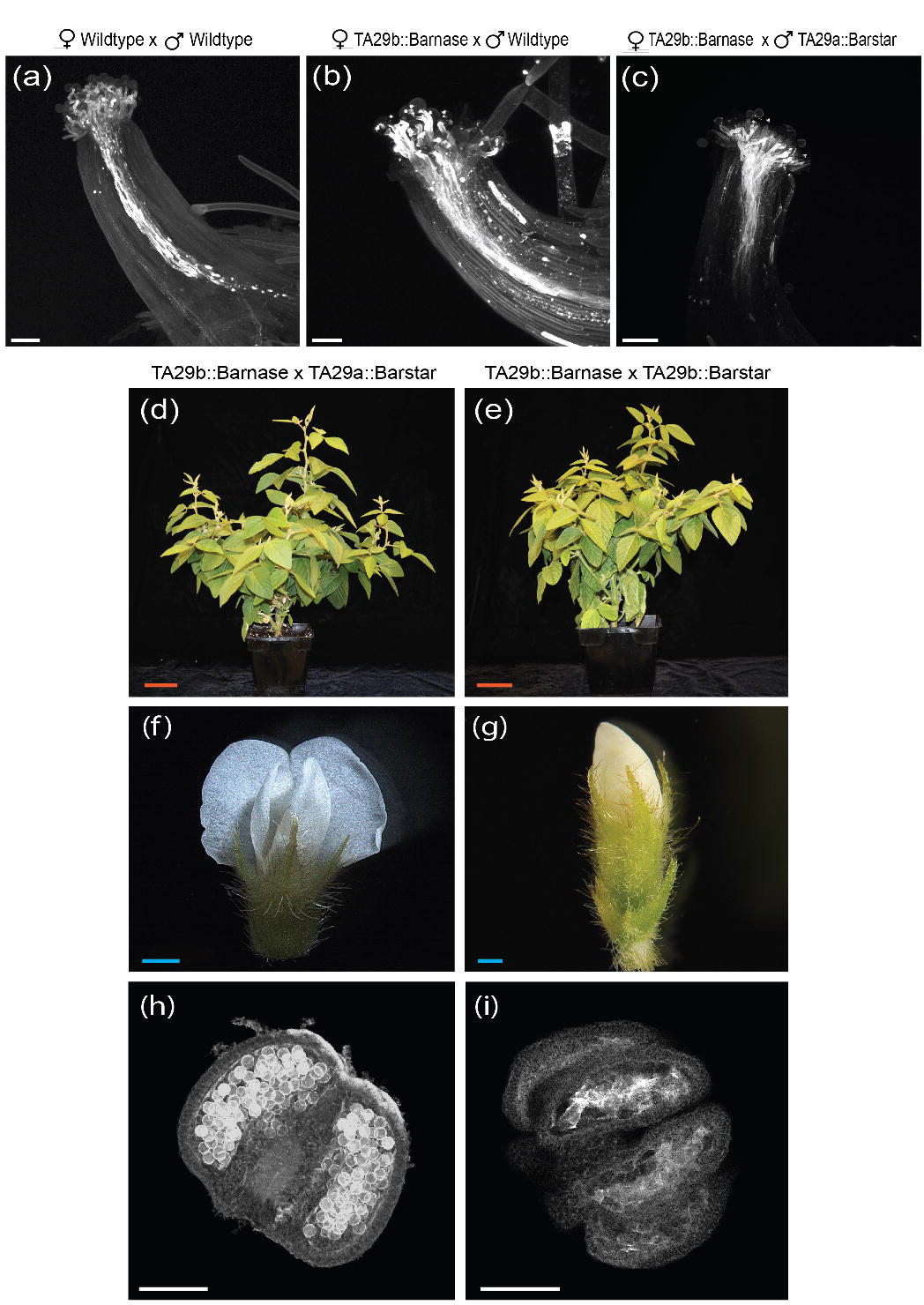
TA29b::Barnase male-sterile carpels are receptive towards pollination and restore fertility to F1 generation when crossed with TA29a::Barstar. (a-c) Aniline blue images capture pollen tube growth through the stile of receptive carpels. (d,e) Plants were grown in growth chamber conditions and imaged 90 days post-germination. (f,g) Flower buds were imaged at peak reproductive maturity. (h,i) PI-stained anthers were dissected from flowers and imaged with a confocal microscope. Scale bars for the following figure are as follows: the white bar is 100μm; the orange bar is 100mm; the blue bar is 1mm.

As mentioned above, we constructed two Barstar rescue lines to test whether a higher dosage of Barstar relative to Barnase is sufficient to fully rescue male fertility in F1 hybrids. We crossed three different TA29a::Barstar and two different TA29b::Barstar lines onto TA29b::Barnase plants, genotyped F1 seedlings for the presence of Barnase and Barstar transgenes (Table S1), and examined male fertility in these F1 hybrids. In line with our findings for the parental transgenic lines (TA29b::Barnase, TA29a::Barstar, and TA29b::Barstar), the presence of both transgenes did not affect overall vegetative growth and morphology (Figure 3d,e).

At reproductive maturity, we noticed that, like the TA29b::Barnase parental lines, all of the TA29b::Barnase x TA29b::Barstar F1 flowers were cleistogamous (i.e. closed at maturity), while TA29b::Barnase x TA29a::Barstar F1 plants developed both cleistogamous and chasmogamous flowers, similar to WT plants (Figure 3f,g). To test whether pollen formation was rescued by crossing with either Barstar construct, we examined PI-stained anthers from our F1 plants using confocal microscopy. While the anther sacs from the TA29b::Barnase x TA29a::Barstar crosses were full of pollen, similar to WT anthers, we found that TA29b::Barnase x TA29b::Barstar anthers formed zero pollen grains (Figure 3h,i). Instead, flowers from TA29b::Barnase x TA29b::Barstar F1s exhibited brightly stained cellular debris inside of the anther sacs, similar to the phenotype that we characterized for TA29b::Barnase flowers (Figure 3i and Figure 2i). Moreover, the only pods that were formed on the TA29b::Barnase x TA29b::Barstar crosses were parthenocarpic, indicating that the plants were completely male-sterile. We, therefore, refer to the crosses with TA29b::Barstar as failed and those with TA29a::Barstar as successful rescues.

Not all TA29b::Barnase x TA29a::Barstar crosses rescued pollen formation to the same degree. We observed one Barnase/Barstar combination, TA29b::Barnase-line 11 x TA29a::Barstar-line 12, that produced clumpy pollen grains that failed to efficiently release from the anthers (Figure S5e). This partial rescue exhibited both cleistogamous and chasmogamous flowers (similar to the full rescue plants); however, aggravated shaking was required to dehisce and release clumpy pollen from the anthers for imaging (Figure S5e). These partial-rescue plants set a mix of reproductive and parthenocarpic pods, however, they formed significantly fewer seeds overall than WT plants (Figure S6). These results indicate that further tuning is needed across the independent Barnase/Barstar lines to ensure consistent rescue phenotypes.

To examine whether successful rescue of pollen formation in TA29b::Barnase x TA29a::Barstar plants can be attributed to higher Barstar expression relative to Barnase, we performed qRT-PCR for Barnase and Barstar on flowers from failed versus successful rescues. We found that F1 plants with rescued fertility exhibited significantly higher Barstar expression relative to Barnase, while failed rescue flowers showed no significant differences between Barnase and Barstar expression (Figure 4a,b). This data supports a model in which higher expression of Barstar relative to Barnase is necessary to rescue pollen formation in F1 hybrids.

**Figure 4:**
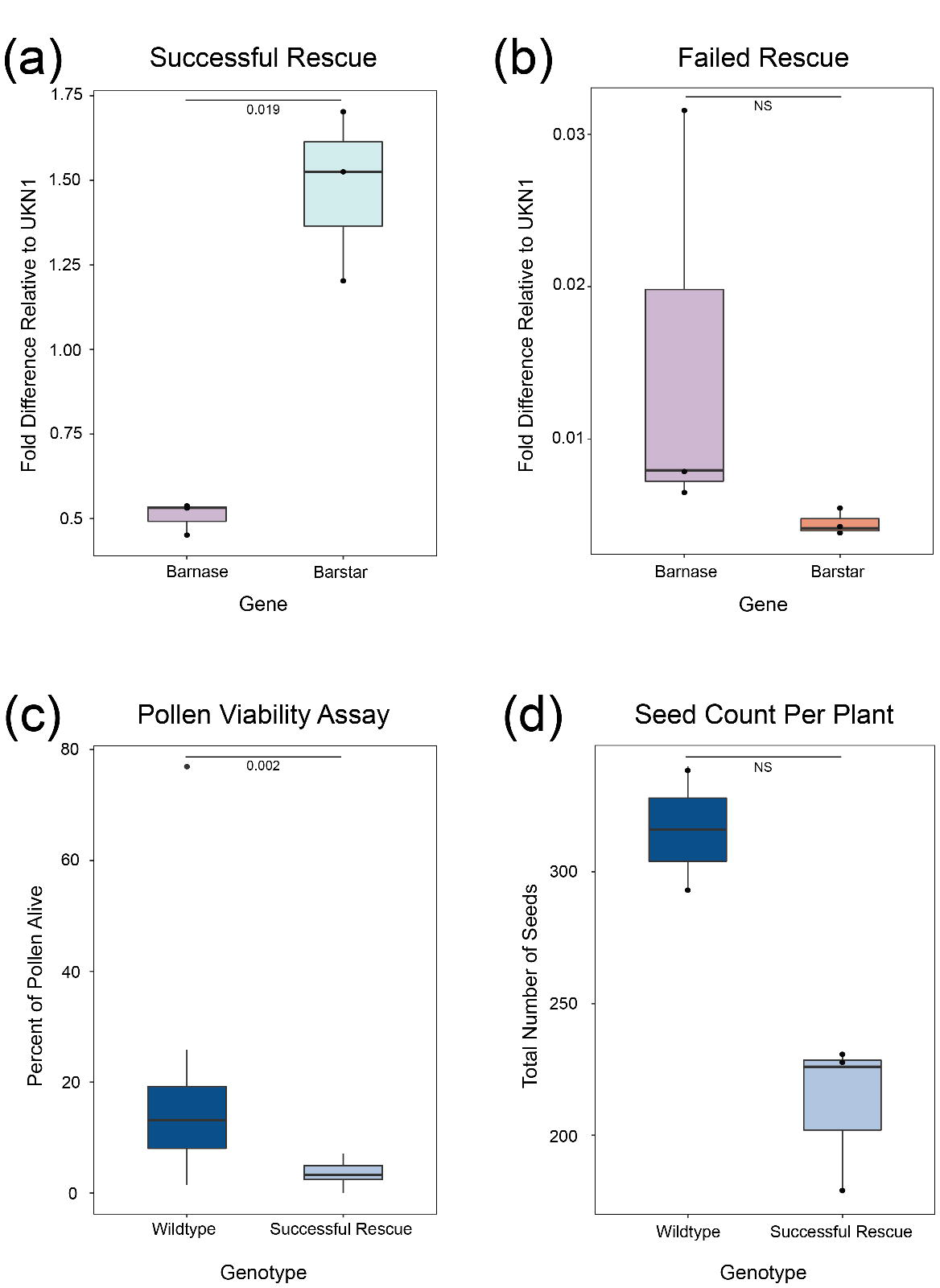
A higher dosage of Barstar contributes to the success of the rescue cross. (a) Boxplot compares qRT-PCR data of Barnase and Barstar genes from the successful rescue line. Barnase and Barstar expressions are normalized relative to the control, UKN1. Bars extending from the boxplot indicate maximums and minimums and the middle line in the box marks the median data point. (b) Boxplot comparing qRT-PCR data of Barnase and Barstar genes from the failed rescue line. (c) A boxplot comparing the percent of pollen alive per image taken during a pollen viability assay. The plot compares wildtype to the successful rescue (n = 36). (d) A boxplot of total seeds collected per plant taken from wildtype and the failed rescue. Statistics were calculated with the student’s t-test.

Next, we tested whether the pollen formed in successful rescue flowers is alive, using a fluorescein diacetate (FDA)/PI pollen viability assay (Figure 4c, Figure S7a-f; (Muhlemann, Younts, and Muday 2018)). We found that all of the successful rescue crosses produced viable pollen; however, overall viability counts for these lines were significantly lower (3.6% viability on average) than in WT flowers (15.7% on average). Moreover, we noticed that WT anthers released substantially more pollen than the successful rescue anthers (Figure S7g). Despite these differences in pollen counts and percent viability, we found that the rescue crosses produced sufficient levels of viable pollen to promote self-fertilization and the formation of reproductive seed pods (Figure 4c,d). When comparing WT plants to successful rescue plants (TA29b::Barnase - line 13 x TA29a::Barstar - line 12) in growth chamber conditions, we found no significant difference in total seed count. These results demonstrate that crossing with TA29a::Barstar can restore reproductive fitness to previously sterile plants containing TA29b::Barnase (Figure 4d).

## Discussion

Here, we demonstrate that Barnase and Barstar can be effectively used to create an obligate outcrossing breeding system in soybean. Notably, we show that a higher dosage of Barstar relative to Barnase is required to produce successful male rescue in the F1 generation (Figure 5). Barnase is a potent ribonuclease; when expressed in the tapetum it destroys this cell layer and inhibits microspore maturation. In this paper, we show that cytotoxic ablation of the tapetum using Barnase is sufficient to produce complete male sterility (Figure 5a). We also demonstrate that male fertility can be rescued in a dosage-dependent manner by expressing the Barnase-inhibitor, Barstar. While an equal dosage of Barstar to Barnase failed to rescue fertility (Figure 5b), when Barstar was expressed at significantly higher levels than Barnase, we were able to recover self-fertile F1 hybrids (Figure 5c).

**Figure 5:**
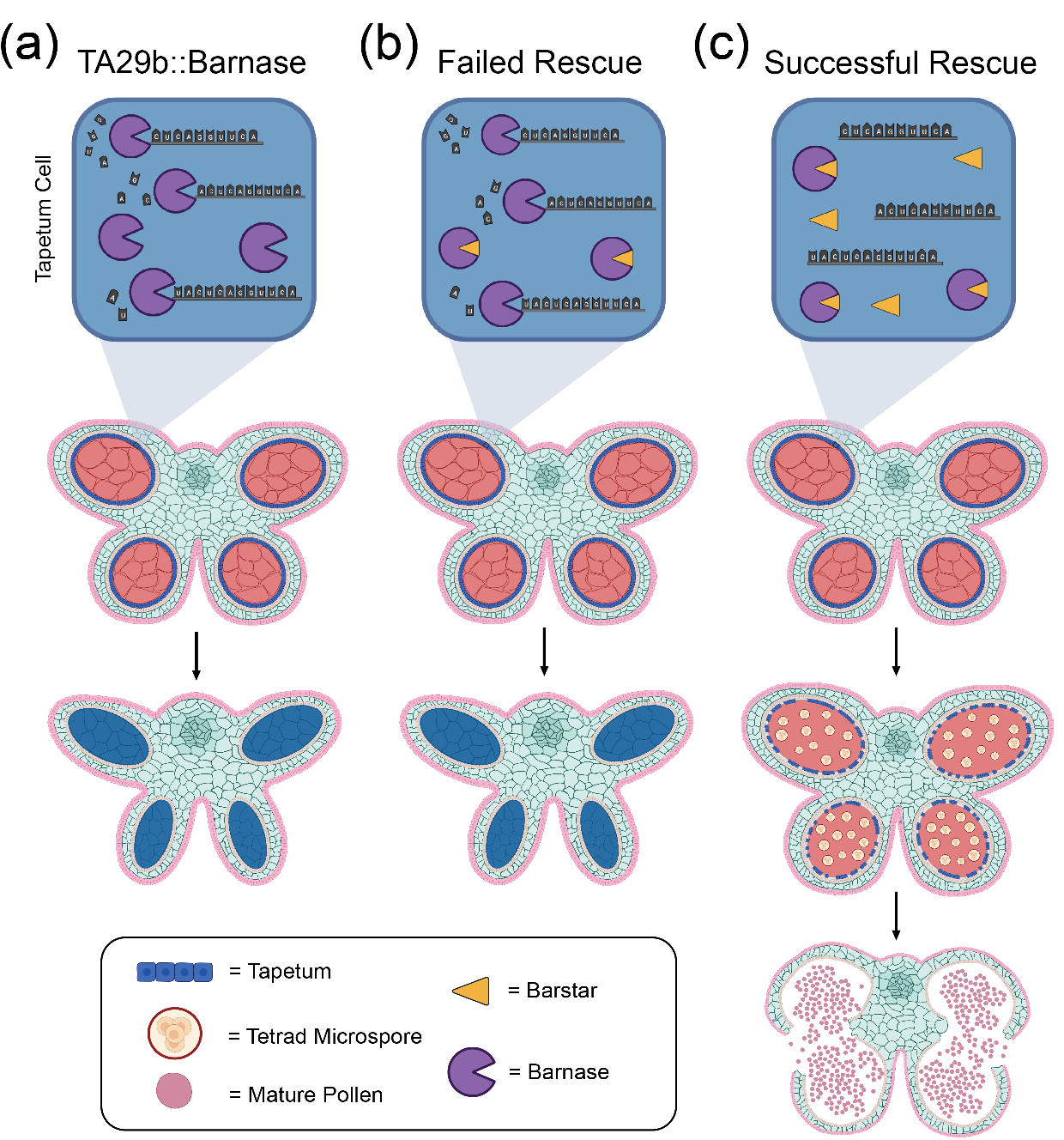
A diagram representation of the effects of TA29b::Barnase, TA29b::Barnase x TA29b::Barstar (failed rescue), and TA29b::Barnase x TA29a::Barstar (successful rescue), respectively. Top row (a-c) depicts Barnase, Barstar, and mRNA interaction inside the nucleus of a single cell in the tapetum. The tapetal cells are enlarged from a digital drawing of soybean anthers containing four pollen sacs. The diagram outlines the development of the anthers under the influence of Barnase and Barstar expression. (c) The successful rescue mimics wildtype anther growth as it progresses from early anther development, microsporogenesis, and anther maturity. (a,b) Pollen development is halted prematurely when Barnase enzymes are uninhibited and the tapetum is destroyed. This figure was created with BioRender.com.

Due to the potent cytotoxic nature of Barnase, a small fraction of uninhibited ribonuclease is sufficient to induce male sterility. In our model, the dosage of Barstar inhibitor must exceed that of the ribonuclease to ensure complete inactivation of Barnase, restoring male fertility (Figure 5c). Even with this successful rescue, we observed some TA29b::Barnase x TA29a::Barstar F1 hybrids that only exhibited partial rescue phenotypes with reduced fertility. These partial rescues indicate that further optimization of the relative dosage of Barnase to Barstar is needed to ensure full rescue and heightened seed set in the F1 hybrids. Tuning of Barnase and Barstar dosage could be achieved by modifying the relative strength of the TA29 promoters through the addition of *cis*-regulatory elements (Biłas et al. 2016) or identifying new tapetum-specific promoters that can be used to regulate Barnase/Barstar expression. Relative to other major crops, reproductive development in soybean is insufficiently studied, in our opinion. Soybean currently lacks key enabling resources such as a detailed floral organ expression atlas, and cell type-specific expression profiling. Such resources are essential for achieving a hybrid breeding system through optimized reproductive rewiring.

To fully realize the potential for hybrid breeding, programmed male sterility/male rescue is only part of the solution; we would also need to recruit insect vectors to facilitate outcrossing. Pollinator ecology studies clearly indicate that honeybees are the primary pollinators in soybean fields (Blettler, Fagúndez, and Caviglia 2018; Delaplane, Mayer, and Mayer 2000). These studies show that a mutually beneficial interaction exists between bees and soybean fields. Soybean fields provide a source of nectar in midwestern agricultural landscapes (Lin et al. 2022), and in the reverse direction, significant increases in crop yield are associated with the presence of nearby apiaries (Blettler, Fagúndez, and Caviglia 2018; Eric H. Erickson 1975; E. H. Erickson et al. 1978; Garibaldi et al. 2021). This work strongly indicates that honeybees could serve as an ecologically beneficial choice for facilitating soybean outcrossing. Building a better understanding of the floral traits that impact honeybee preferences in soybean is essential to effectively incorporate honeybees as vectors for hybrid breeding.

Furthermore, we currently lack a clear framework for predicting strong heterotic combinations in soybean, which precludes our ability to project the overall benefits that could be derived from applying hybrid breeding to this major crop. This is in large part due to the technical challenges of generating hybrid seeds for this species. Indeed, almost all hybrid soybean seed is generated by making crosses by hand, which means that research on heterosis in this crop has been restricted to a small number of crosses with low sample sizes (Burton and Brownie 2006; Palmer et al. 2001). Though information is limited, existing studies show that particular combinations of inbred parents can lead to significant yield boosts in the F1 generation (Burton and Brownie 2006; Palmer et al. 2001). However, to date, these studies are limited to narrow geographic regions due to the challenges of generating hybrid seeds. Given that soybean is grown globally across a broad range of environments (Ates and Bukowski 2023b), current knowledge on hybrid vigor in soybean does not necessarily apply to this crop’s expansive geographic range, emphasizing the need for efficient hybrid breeding (E. H. Erickson 1975). Obligate outcrossing with the Barnase/Barstar lines that we present in this paper provides a new resource that can be used to amplify hybrid seed sets, enabling large-scale trials for heterosis in this major crop.

Hybrid breeding in soybean has the potential to increase the productivity of one of the most planted and consumed crops in the Americas, yet it has remained largely unexplored. This is in part due to the limitations of current approaches, which have failed to produce reliable obligate outcrossing in soybean. The work we present in this paper provides a key enabling technology to produce obligate outcrossing in soybean.

## Materials and Methods

### Construct Design and Synthesis

Constructs for tapetum-specific expression of Barnase/Barstar proteins were designed based on sequences from *Bacillus amyloliquefaciens.* Two soybean paralogs for TA29 were identified using a BLAST search with the Arabidopsis promoter for TA29 (At5g07230) as a query sequence. This query uncovered TA29a (Glyma.09G144000) and TA29b (Glyma.16G197100). Expression data for the two soybean TA29 paralogs was downloaded from the JGI Plant Gene Atlas (Sreedasyam et al. 2022; Valliyodan et al. 2019). To ensure the full rescue of Barnase with the Barstar rescue lines, a relatively weaker expressing TA29b::Barnase construct, and two versions of Barstar constructs, TA29a::Barstar, and TA29b::Barstar were generated (Figure 1a-c, Files S1-3). In addition, to confirm that TA29a/TA29b are expressed in tapetal cells, a dual reporter construct expressing TA29a::td-Tomato and TA29b::ZsGreen in opposite directions with an insulator (Transformation booster sequence (TBS) from petunia) between the two reporters was generated (Figure 1d, File S4). All construct sequences can be found in Files S1-4. Cloning was achieved by the Wisconsin Crop Innovation Center (WCIC) using the Golden Gate MoClo Plant Tool Kit and final vector backbone pAGM4673 provided in the kit (Wisconsin Crop Innovation Center, Madison, WI) (Engler et al. 2014).

### Plant Transformation

Transgenic soybean lines were generated at the WCIC using a proprietary protocol that involves *Agrobacterium tumefaciens*-mediated transformation of cultured shoot meristems. Positive transformants were selected based on kanamycin resistance and confirmed with transgene-specific primers. At least 8 independent transformants were generated for each construct.

### Genotyping Transformants

All T1 plants used in this study were genotyped for transgene inheritance using primers targeted to the aadA1 antibiotic resistance gene, which was present in all of the constructs, and construct-specific targets: Barnase, Barstar, ZsGreen, and td-Tomato (Primers, PCR conditions, and construct details are in Table S1). Genotyping PCRs were performed using Promega Hotstart Green MasterMix following the recommended conditions from the manufacturer (Promega, Madison, WI, U.S.A.).

### Growth chamber and greenhouse conditions

Plants were grown in a greenhouse (daytime temp: 23.9°C in heating and 26.7°C in cooling; nighttime temp: 20°C in heating and 22.8°C in cooling, light for 14 hours) and growth chamber (23°C, 100% light for 16 hours, R/H at 60%) facilities at Cornell University. Plants in Figure S4 were grown in greenhouse conditions at the Donald Danforth Plant Science Center (daytime temp: 25°C day/ 23°C night; 35% minimum humidity; light for 14 hours (supplemental light when sunlight is below 400 W/m^2^ and shade curtain pulls down to 50% when sunlight is over 900 W/m^2^ and pulls down to 100% when sunlight is over 1000 W/m^2^).

### Soybean crosses with TA29b::Barnase lines

Mature TA29b::Barnase flowers were identified based on the emergence of petals from under the sepal whorl. At this stage in normal flower development, the pollen is viable and the stigma is receptive to pollination (Talukdar and Shivakumar 2012). To uncover the stigma, sepals and petals were manually removed from the TA29b::Barnase flowers, and anthers from pollen donors were rubbed directly onto the stigma surface. To increase the probability of fertilization, multiple crosses were made from the same pollen donor onto each stigma.

### Staining and Imaging Samples

To obtain light microscopy images of whole flowers and individual whorls, soybean flowers were dissected under a Leica M205 FCA fluorescent stereo microscope and imaged with a DMC6200 camera.

For propidium iodide staining, samples were fixed in fresh FAA (50% Ethanol, 10%, 37% Formaldehyde, 5% glacial acetic acid) in small glass vials. Samples were vacuum infiltrated until the majority of the samples sunk to the bottom of the vial. The flower samples were transferred to cuvettes and placed in closed cups containing fresh FAA and incubated overnight at 4°C. The samples were transferred to 50% ethanol at room temperature, and then gradually dehydrated through an ethanol series for 20 minutes each, followed by incubation in 2 x 100% ethanol for an hour each, and gradually rehydrated (20 minutes each) to 100% water. Rehydrated samples were rinsed with sterile, autoclaved water twice, and then stained with Propidium Iodide (2mL of 1mg/ml PI stock in 100mL + 0.02% DMSO) for one hour. Samples were rinsed 2 x 20 minutes in water, gradually dehydrated to 100% ethanol, and then transferred into 50:50 EtOH:Methyl Salicylate for 2 hours, followed by 100% methyl salicylate for 1-2 weeks at 4°C for clearing. Completely cleared samples were determined based on their translucent appearance and imaged on an LSM880 Confocal multiphoton inverted - i880 Zeiss microscope. Samples were imaged using either 514nm or 561nm excitation, collecting in the 566-718nm emission range, with laser power ranging from 0.02-0.024%.

To image TA29a::td-Tomato/TA29b::ZsGreen dual reporter lines, flowers were cleared following a previously published ClearSee protocol to reduce background autofluorescence (Kurihara et al. 2015). Young soybean flower buds (approximately 3-4 mm in length) were fixed with 4% (w/v) paraformaldehyde for 2 hours in PBS under a vacuum at room temperature. The fixed tissue was washed twice with PBS for 1 minute each and cleared with ClearSee solution (10% Xylitol (w/v), 15% Sodium deoxycholate (w/v), 25% Urea (w/v)) at room temperature for 4 weeks under gentle shaking (120 rpm). The clearing solution was changed every other day. After confirming that the tissue was translucent, young buds were imaged using an LSM880 Confocal multiphoton inverted - i880 Zeiss microscope. ZsGreen fluorescence was imaged with a 488nm laser, collecting in the 493-556nm emission range, with laser power at 0.025%. td-Tomato fluorescence was imaged with a 561nm laser, collecting in the 566-691nm emission range, with laser power at 0.02%. The FIJI software was used to merge image data from td-Tomato and ZsGreen (Schindelin et al. 2012).

### Pollen viability and pollen tube growth assays

Pollen viability for Barstar, WT, and Barnase/Barstar rescue crosses was quantified using a previously published protocol (Muhlemann, Younts, and Muday 2018). Briefly, anthers were dissected during anthesis and placed in a 15-mL conical tube. Pollen was released from the anthers by vortexing. The released pollen was resuspended in pollen viability solution [PVS, 290 mM sucrose, 1.27 mM Ca(NO3)2, 0.16 mM boric acid, 1 mM KNO3) containing 0.001% (wt/vol) fluorescein diacetate, and 10 μM propidium iodide (stock solutions: 1% (wt/vol) fluorescein diacetate in acetone; 2 mM PI in water]. The pollen was stained for 15 min at 28°C and then centrifuged. The PVS containing FDA and PI was replaced with PVS alone and the stained pollen was placed on a microscope slide, and then imaged by confocal microscopy on a Zeiss 880 LSCM microscope. The FDA was excited with a 488nm laser and its signal was collected at 493–584nm. The PI was excited with a 561nm laser and its signal was collected at 584– 718nm. Pollen was hand-counted within the FIJI software using the cell counter plugin (Schindelin et al. 2012).

To highlight pollen tube development, a modified protocol for using aniline blue staining was followed (Nasrallah, Liu, and Nasrallah 2002). Hand-crossed and self-pollinated flower buds were harvested and fixed in Farmer’s Solution (3:1 100% ethanol: glacial acetic acid) 24 hrs post-pollination. Buds were incubated at room temperature for 5 minutes and then transferred into 5 M NaOH. Samples were incubated at 65°C for 30 minutes to soften the plant tissue, the buds were carefully rinsed three times in distilled water, and then placed in decolorized aniline blue solution [0.1% (w/v) aniline blue dissolved in 0.1 M K_3_PO_4_ and decolorized by incubating overnight at 37°C in the dark]. After 30 minutes, the carpel was dissected, the trichomes were removed, and the sample was arranged in a drop of aniline blue solution on the microscope slide. The cover slip was added, and pressure was gently applied to squash the tissue. Slides were sealed with clear topcoat nail polish to avoid drying, and examined with an LSM880 Confocal multiphoton inverted - i880 Zeiss microscope, exciting with a 405nm diode laser and collecting in the 415-735nm emission range, or on an Olympus upright Metamorph microscope using UV fluorescence and a DAPI filter.

### CropReporter™ Imaging

Plants were imaged over the course of multiple days at late vegetative and flowering stages using the CropReporter™ imaging box (Netherlands Plant Eco-phenotyping Centre (NPEC), Wageningen University & Research and Utrecht University). From a top view, the CropReporter™ collected dFv/Fm, F0, Fm, RGB, NIR, CHL, and dCHL measurements in dark and light-adapted plants to calculate the Anthocyanin-index (Anthocyanin: (R_550_)^−1^-(R_700_)^−1^), the Chlorophyll-index (Chlorophyll: (R_700_)^−1^-(R_NIR_)^−1^), and the Fv/FM (Fv/Fm = Fm - F0/Fm). Fv/Fm was used to determine if a stressor was affecting photosystem II efficiency. The Phenovation Data Analysis software v548 was used to separate the plant from its background and segregate collected data into individual heatmaps for each type of information collected. The heat maps visualized regions of high anthocyanin pigmentation, high photosystem II efficiency, and overall plant stress.

### Molecular Characterization

For genotyping, DNA was extracted using a modified protocol (King et al. 2014). 400 ul of Edward’s Buffer [40% (v/v) 5 M NaCl and 60% (v/v) extraction buffer (200 mM Tris/HCL pH 7.5, 250 mM NaCl, 25 mM EDTA, 0.5% SDS)] was added to a 1.5 ml microcentrifuge tube containing a young trifoliate leaf, ground up with a micro pestle and incubated at 60℃ for 30 minutes. After incubation, samples were centrifuged at 9,000 x g for 5 minutes. 270ul of the supernatant was transferred to a new tube and gently mixed with 270ul of isopropanol. The new tubes were incubated at −20℃ for 15 minutes, followed by centrifugation at full speed for 5 minutes. Following centrifugation, the supernatant was removed, and the pellet was washed with 70% EtOH, centrifuged at full speed for 5 minutes, and decanted. Residual EtOH at the bottom of the tube was removed via micropipette and the tube was left open to dry at 60℃ for 10 minutes. Once all the ethanol was removed, sterile water was added and the tube was incubated at 60℃ for a final 30 minutes.

### Quantifying Barnase/Barstar Expression

Gene expression was quantified using qRT-PCR. RNA was extracted from young soybean buds using TRIzol (Ambion, Austin, TX, U.S.A), following the manufacturer’s instructions for plant extraction. GlycoBlue™ Blue Coprecipitant (Invitrogen, Waltham, MA, U.S.A) was added to each sample during the RNA precipitation step to facilitate recovery of the RNA pellet. The concentration and quality of the RNA were checked using a DeNovix DS-11 FX+ Spectrophotometer/Fluorometer. RNA samples with 260/280 ratios ranging from 1.7-2.2 were converted to cDNA using QuantiTect™ Reverse Transcriptase (Qiagen, Hilden, Germany). cDNA concentrations were quantified using a nanodrop and diluted to 1-10ng of cDNA, and 1ul of diluted cDNA was added to Power Up™ SYBR Green Master Mix (Applied Biosystems, Waltham, MA, U.S.A) along with gene-specific primers targeting ZsGreen, td-Tomato, Barnase, Barstar, and a housekeeping gene, UKN1 (Table S2). The qRT-PCR housekeeping gene, UKN1 was selected based on a literature review of stable targets used for soybean gene expression experiments (Hu et al. 2009; Jian et al. 2008). Samples were run on a Bio-Rad CFX96 Optics Module with a C1000 Touch Base set at 50℃ for 2 minutes followed by 95℃ for 2 minutes for predenaturation. For 39 cycles, the machine was set at 95℃ for 10 seconds and 60℃ for 30 seconds to denature, anneal, and elongate. Downstream data analysis was performed with an Excel spreadsheet to compute the delta Ct value (normalized to the housekeeping gene, UKN1) and Student’s t-test for significance.

## Acknowledgments

This work was funded by a gift from Dr. Phil Needleman to the Donald Danforth Plant Science Center; an award from the United Soybean Board 1920-152-0131-D; and a Foundation for Food and Agricultural Research New Innovator in Food and Agriculture Research Award FF-NIA21–0000000017. Imaging data was acquired through the Cornell Institute of Biotechnology’s Imaging Facility, with NIH S10OD018516 funding for the shared Zeiss LSM880 confocal/multiphoton microscope.

