## Supplemental Figures for "Introduction of Barnase/Barstar in soybean produces a rescuable male sterility system for hybrid breeding"

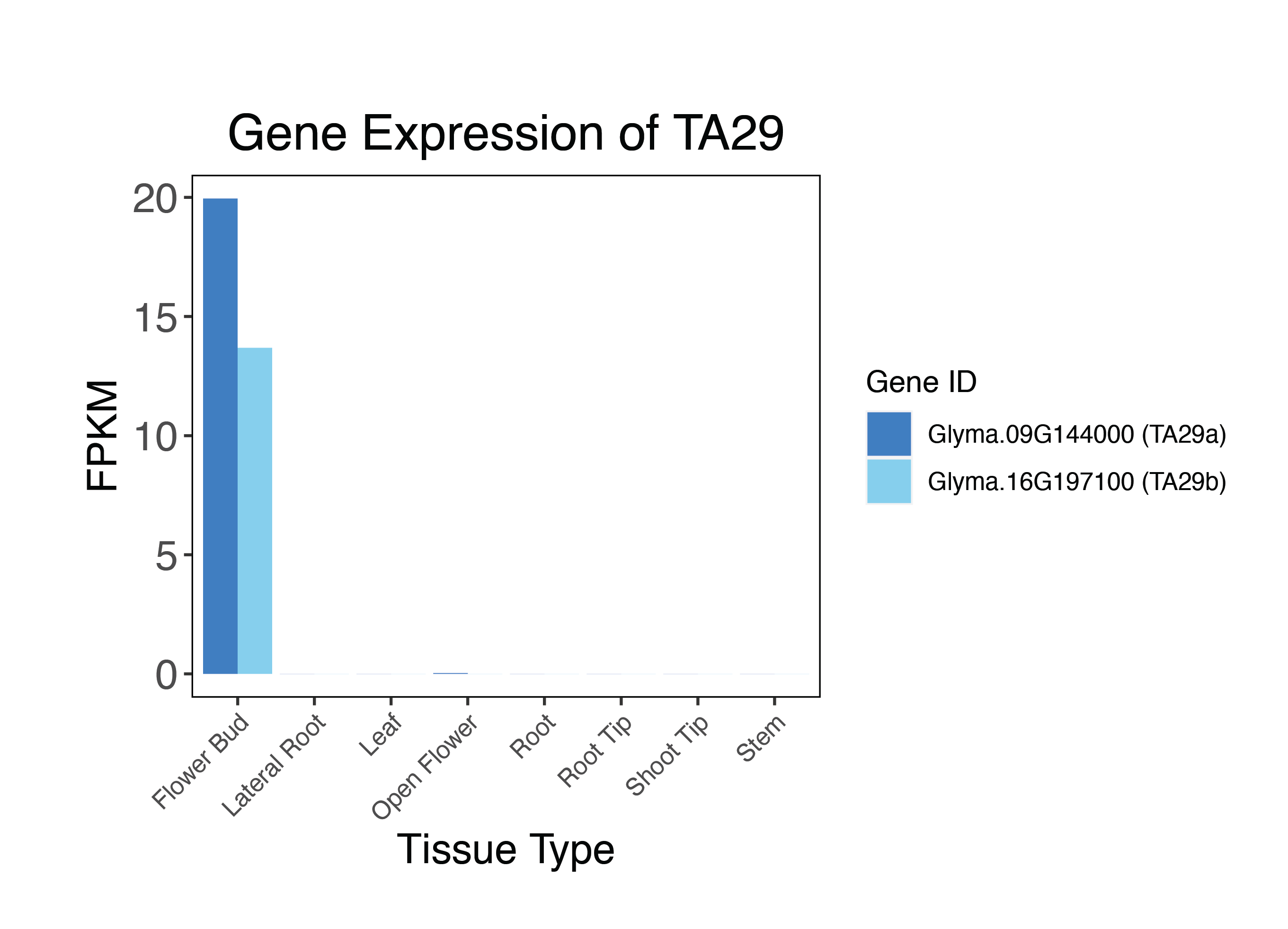

Supplemental Figure 1: Paralogs of TA29 are differentially expressed in flower buds. RNAseq data published on JGI Plant Gene Atlas compares fragments per kilobase of exon per million mapped fragments (FPKM) for TA29a and TA29b.

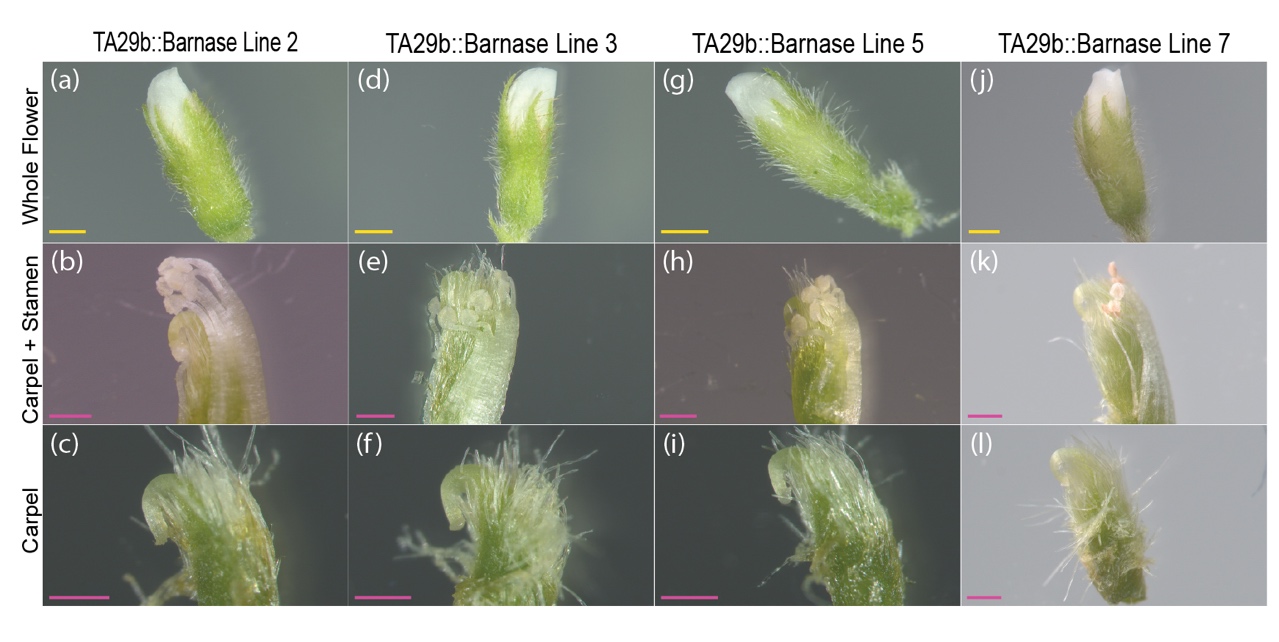

Supplemental Figure 2: TA29b::Barnase transformation events result in cleistogamous flowers, sterile anthers, and curved carpels. (a-l) Flower dissection images of four additional positive transformation events of TA29b::Barnase. The scale bars are as follows: the yellow bar is 1mm and the pink bar is 500μm.

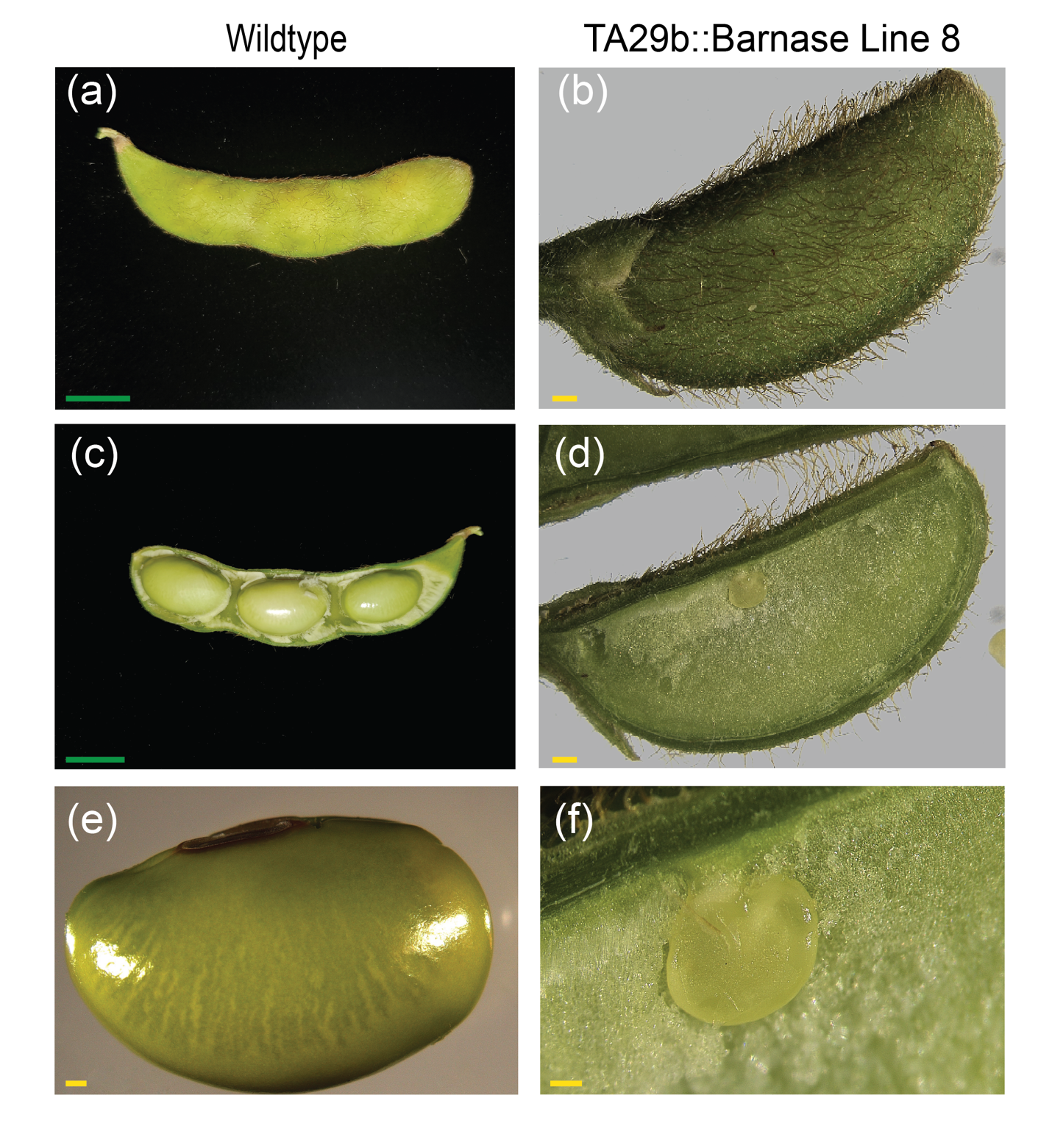

Supplemental Figure 3: Male sterile Barnase lines produce parthenocarpic pods. At maturity, wildtype pods elongate (a,c) and develop seeds (e). Parthenocarpic pods (b,d) remain short and the ovules never develop into embryos (f). Images (b,d-f) were taken on a dissecting microscope. The scale bars are as follows: the green bar is 10mm and the yellow bar is 1mm.

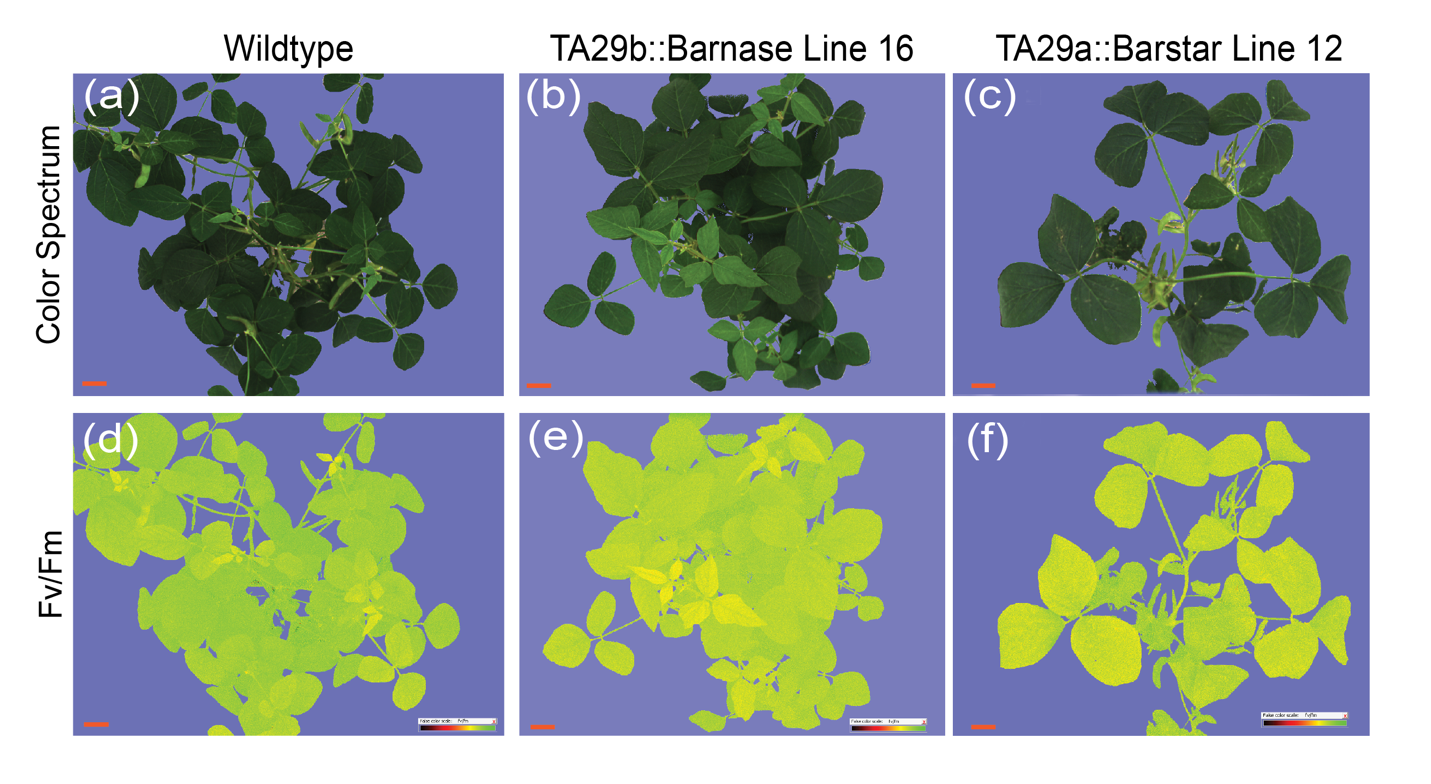

Supplemental Figure 4: No stressors are acting upon TA29b::Barnase and TA29a::Barstar transformation events. Top-view images of wildtype, TA29b::Barnase, and TA29a::Barstar lines were captured with NPEC’s CropReporter™. Color spectrum images (a-c) pair with Fv/Fm images (d-f) to demonstrate no stress is present in the plant by the appearance of bright green on the Fv/Fm scale. Plants were grown under greenhouse conditions at the Danforth Plant Science Center. The orange scale bar is 100mm.

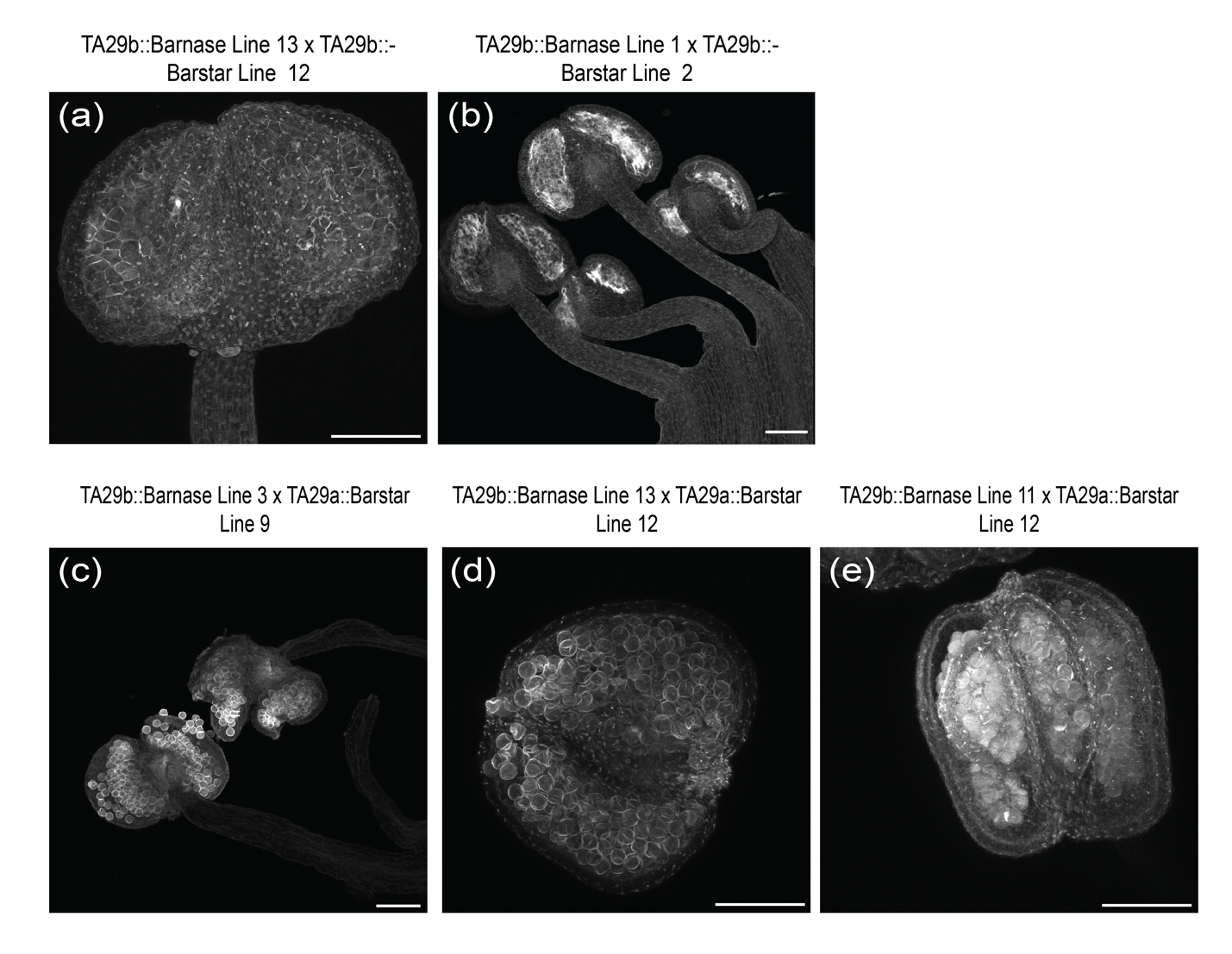

Supplemental Figure 5: TA29b::Barstar is unable to rescue fertility in the F1 generation. (a,b) Additional PI-stained anther images captured by confocal microscopy of failed rescue lines display consistent male sterility regardless of TA29b::Barnase transformation event. (c,d) PI-stained images of successfully rescued anthers highlight the existence of pollen granules in varying Barnase/Barstar transformation line pairings. (e) PI-stained anther of partial rescue develops pollen which clumps together and away from the anther wall. The white scale bar is 100μm.

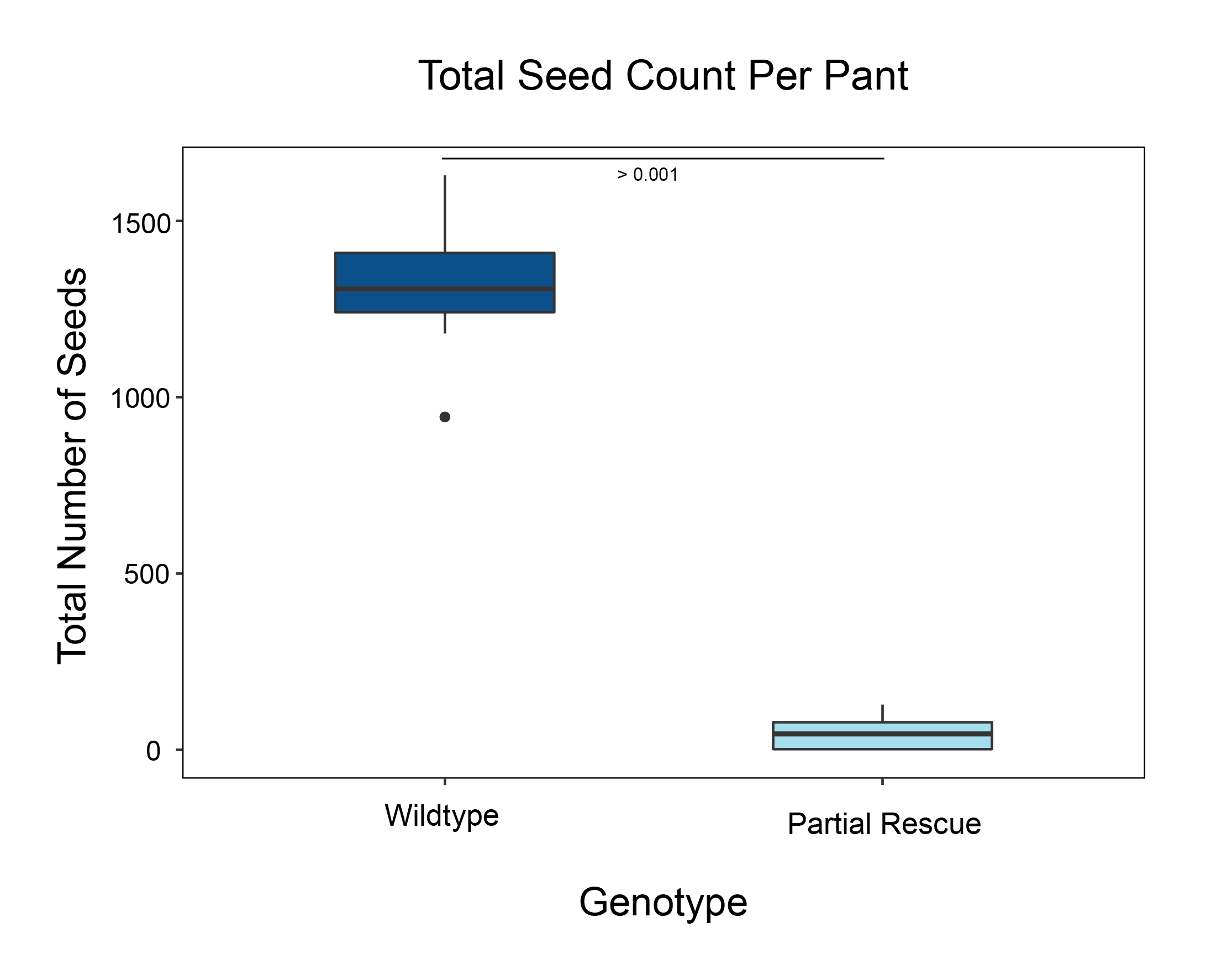

Supplemental Figure 6: Partial rescue produces significantly less seeds compared to wildtype. The boxplot graphs the significant increase in number of seeds per plant for wildtype compared to partial rescues grown in greenhouse growing conditions (n=22). Statistics were done with a two-tailed Student’s t-test.

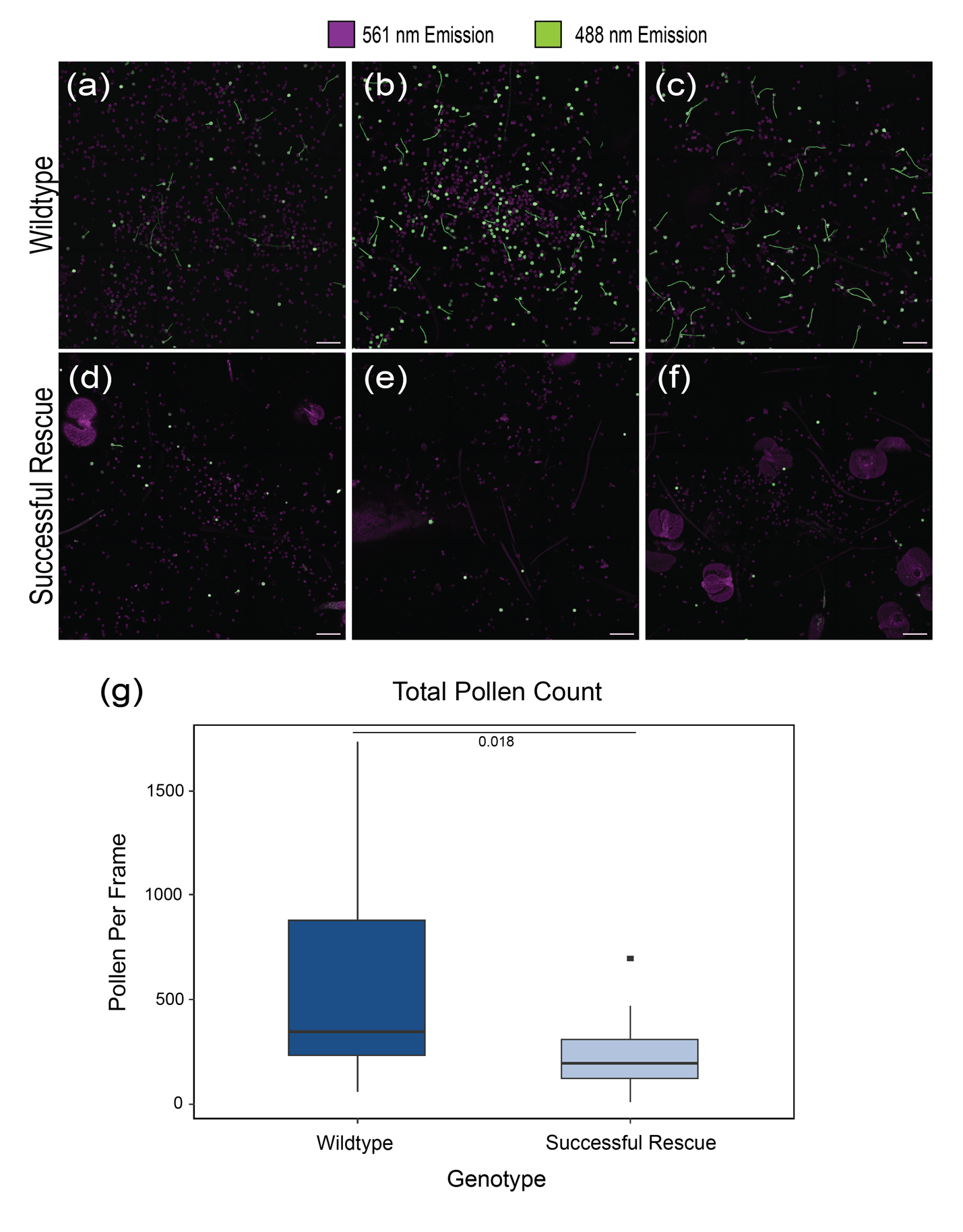

Supplemental Figure 7: Successful rescue lines produces significantly less pollen compared to wildtype. The pollen viability assay stains living cells with FDA, emitting green fluorescence at 488nm emission, and stains dead cells with PI, emitting red fluorescence at 561nm emission by laser lines on the confocal microscope. In this figure, the red color has been converted to magenta using the Fiji Software system. Wildtype assay images (a-c) have a significantly higher number of pollen granules per 9-tile frame than successful rescue assay images (d-f). This data is reflected in boxplot (g, n=37). Statistics were done using a two-tailed Student’s t-test. The light pink scale bar is 200μm.

| Primer Name | Primer Sequence | Annealing Temperature |
| --- | --- | --- |
| aadA1_Forward | CTCCATCAACAGCAGATCCA | 63℃ |
| aadA1_Reverse | CGCACAATCCCACTATCCTT | 63℃ |
| Barstar_Forward | GAAAGAACTCGCTCTGCCAG | 63℃ |
| Barstar_Reverse | TGCAAAACACTTTCAGCTCCA | 63℃ |
| Barnase_Forward | TTCGACGGTGTTGCAGACTA | 63℃ |
| Barnase_Reverse | CGGTTACTAAAGATGTCGCCC | 63℃ |
| ZsGreen_Forward | CCAGTTATCGGTCATCTTCTTC | 60℃ |
| ZsGreen_Reverse | TACCACGAGTCCAAGTTCTAC | 60℃ |
| td-Tomato_Forward | TGGCTACTACTACGTCGATAC | 60℃ |
| td-Tomato_Reverse | GTGACGTCCTTCTGATCTTTC | 60℃ |

Supplemental Table 1: A table of the primers used for genotyping transgenic lines.

| qRT-PCR Primer Name | Primer Sequence | Annealing Temp | Target Gene |
| --- | --- | --- | --- |
| Barnase_qRT-PCR_Forward | GGGTAGCTTCAAAGGGTAAC | 60℃ | N/A |
| Barnase_qRT-PCR_Reverse | TCCAGGCAATTTACCTTCAC | 60℃ | N/A |
| Barstar_qRT-PCR_Forward | TGGAGAACAGATCAGGAGTATC | 60℃ | N/A |
| Barstar_qRT-PCR_Reverse | CAGTCCCAAAGAGCATCAAG | 60℃ | N/A |
| ZsGreen_qRT-PCR_Forward | CCAGTTATCGGTCATCTTCTTC | 60℃ | N/A |
| ZsGreen_qRT-PCR_Reverse | TACCACGAGTCCAAGTTCTAC | 60℃ | N/A |
| td-Tomato_qRT-PCR_Forward | TGGCTACTACTACGTCGATAC | 60℃ | N/A |
| td-Tomato_qRT-PCR_Reverse | GTGACGTCCTTCTGATCTTTC | 60℃ | N/A |
| UKN1_qRT-PCR_Forward | CTGGATCTGGCACTCTTATTC | 60℃ | Glyma.12g02310.1 |
| UKN1_qRT-PCR_Reverse | GAGCTGAGTTTGCATGTTTG | 60℃ | Glyma.12g02310.1 |
| FLD/LSD1_ qRT-PCR_Forward | CTCCTGAGGTTTGGGTTTAAG | 60℃ | Glyma.02g159100.1 |
| FLD/LSD_ qRT-PCR_Reverse | CAGCAGCACACATCCTATTC | 60℃ | Glyma.02g159100.1 |
| CYP2_ qRT-PCR_Forward | CGGATCTCAGTTCTTCATCTG | 60℃ | Glyma.12g024700.1 |
| CYP2_ qRT-PCR_Reverse | TGGATCCGACCTTCTCTATC | 60℃ | Glyma.12g024700.1 |

Supplemental Table 2: A table listing primers used for qRT-PCR.

[Supplemental File 1](https://drive.google.com/file/d/1sRVI6IilgJS724lx3xXiZvHZl5V9OWIF/view?usp=share_link) - TA29b::Barnase Construct

[Supplemental File 2](https://drive.google.com/file/d/1Ex_wVZiS4IDgZ-W9Z-mAFyzEYXOCGEyX/view?usp=share_link) - TA29a::Barstar Construct

[Supplemental File 3](https://drive.google.com/file/d/1s9wL0Xu6RpRPC9OxjRx6he-_H4KOZZ9q/view?usp=share_link) - TA29b::Barstar Construct

[Supplemental File 4](https://drive.google.com/file/d/1oY7nK-8rWIuapCtJZgbrX6w2V8SzyqEN/view?usp=share_link) - TA29a/b Control Construct
